# Murine cytomegalovirus downregulates ERAAP & induces an unconventional T cell response to self

**DOI:** 10.1101/2022.08.23.504566

**Authors:** Kristina M. Geiger, Michael Manoharan, Rachel Coombs, Chan-su Park, Angus Y. Lee, Nilabh Shastri, Ellen A. Robey, Laurent Coscoy

**Author notes:** Lead contacts (Ellen A. Robey, Laurent Coscoy).

## Abstract

The endoplasmic reticulum aminopeptidase associated with antigen presentation (ERAAP) plays a crucial role in shaping the peptide-MHC I repertoire and maintaining immune surveillance. While murine cytomegalovirus (MCMV) has multiple strategies for manipulating the antigen processing pathway to evade immune responses, the host has also developed ways to counter viral immune evasion. In this study, we found that MCMV modulates ERAAP and induces an IFN-γ producing CD8^+^ T cell effector response that targets uninfected ERAAP deficient cells. We also observed that ERAAP downregulation during infection led to presentation of the self-peptide FL9 on non-classical Qa-1b, thereby eliciting Qa-1b restricted QFL T cells to proliferate in the liver and spleen of infected mice. QFL T cells upregulated effector markers upon MCMV infection and were sufficient to reduce viral load after transfer to immunodeficient mice. Our study highlights the consequences of ERAAP dysfunction during viral infection and provides potential targets for antiviral therapies.

## Introduction

Viral immune surveillance relies on the antigen processing pathway to generate viral-derived peptides for presentation on Major Histocompatibility Complex (MHC) I molecules on the surface of infected cells for recognition by cytotoxic CD8^+^ T cells (CTL) (Blum et al., 2013). Peptide and MHC (pMHC) complexes are generated through a series of coordinated steps in which both microbial and self-proteins are initially processed in the cytosol by the proteasome (Kloetzel, 2004; Rock et al., 2002) then transported into the endoplasmic reticulum (ER) through the transporter associated with antigen processing (TAP) (Van Kaer, 2002; Rock et al., 2002) and finally trimmed in the ER by ERAAP to the optimal length of 8-10 amino acids for MHC I binding (Kanaseki et al., 2006; Shastri et al., 2002; Yewdell et al., 2003). To escape CTL detection, many viruses, including human cytomegalovirus (HCMV) and its mouse homolog murine cytomegalovirus (MCMV) have evolved multiple strategies for targeting MHC I and the antigen processing pathway (Halenius et al., 2015; Kavanagh et al., 2001a). MCMV encodes viral proteins m4, m6, and m152 that target MHC class I through multiple mechanisms including retaining MHC I at the ER-Golgi intermediate compartment and redirecting MHC I for lysosomal degradation (Doom and Hill, 2008; Kavanagh et al., 2001b; Reusch et al., 1999; Wagner et al., 2002; Ziegler et al., 1997). In addition to directly targeting MHC I (Halenius et al., 2011; Jackson et al., 2011), HCMV also targets ERAAP (ERAP1) mRNA through both miR-UL112-5p and miR-US4-1 to inhibit processing of HCMV peptides (Kim et al., 2012; Romania et al., 2017). The impact of viral-mediated ERAAP downregulation on CD8^+^ T cell responses, and whether MCMV also targets ERAAP, remain unclear.

In the absence of infection, loss of ERAAP leads to an altered display of MHC 1 bound self-peptides (Nagarajan and Shastri, 2013; Nagarajan et al., 2016). As a result, immunization of wild type mice with ERAAP deficient cells induces a CTL response against these novel pMHC complexes (Nagarajan et al., 2012). Interestingly, a substantial proportion of this response consists of T cells directed against the non-classical MHC Ib molecule Qa-1b presenting a peptide (FL9) derived from broadly expressed proteins, Fam49A/B (termed “QFL” T cells). In addition to using a non-classical MHC Ib restricting molecule, QFL T cells have other characteristics of unconventional T cells, including the predominant use of a fixed TCR alpha chain and an antigen experienced (CD44+) phenotype in naïve mice (D’Souza et al., 2019; Guan et al., 2017). It has been suggested that QFL T cells may monitor cells for ERAAP dysfunction (Nagarajan et al., 2012), but whether QFL T cells play a role during infection remains unknown.

Qa-1b, along with its functional homologs HLA-E in humans and Mamu-E in rhesus macaques (collectively called MHC-E), are members of a large family of non-classical MHC Ib molecules. Unlike classical MHC Ia molecules, which are polymorphic, bind diverse sets of peptides, and are recognized by diverse αβ T-cell receptors (TCR) on CD8^+^ T cells, non-classical MHC Ib molecules are non-polymorphic, present a limited set of peptides and non-peptidic ligands, and in some cases can be recognized by NK cell receptors as well as some semi-invariant TCRs (Allen and Hogan, 2013; Rodgers and Cook, 2005). In healthy cells, Qa-1b and HLA-E predominantly present conserved leader peptides from MHC Ia molecules (termed Qdm in mouse and VL9 in human) and serve as ligands for NK receptors (e.g. CD94-NKG2A) (Kurepa et al., 1998; Sharpe et al., 2019; Vance et al., 1999). Subsequently it was shown that Qa-1b and HLA-E molecules can also present self and microbial peptides to CD8^+^ T cells (Lo et al., 1999; Oliveira et al., 2010; Sell et al., 2015; Shang et al., 2016). Recent interest in MHC-E has been stimulated by a rhesus macaque CMV (Rh-CMV) vectored vaccine that elicits a broad and protective MHC-E restricted CD8^+^ T cell response against simian immunodeficiency virus (SIV) infection (Hansen et al., 2011, 2013a, 2013b, 2016, 2019; Malouli et al., 2021; Verweij et al., 2021; Yang et al., 2021). Because of the limited polymorphism of MHC-E, together with the promising results from the SIV vaccine, unconventional MHC-E restricted T cells show great potential in vaccine development. However, our limited understanding how MHC-E restricted T cell responses are generated and how they provide protection hampers our ability to realize the clinical potential of these responses.

MHC-E restricted CD8^+^ T cells have been described in both HCMV and MCMV infection (Anderson et al., 2019; Jouand et al., 2018). However, whether these T cell responses are shaped by and respond to ERAAP downregulation during infection is not known. In this study, we find that ERAAP protein levels are downregulated in MCMV infected cells, resulting in a strong immune response that targeted uninfected ERAAP knockout cells. MCMV infection *in vivo* stimulated the expansion and effector differentiation of MHC-E restricted QFL T cells, and QFL T cells were protective in MCMV infected Rag2/γc knockout mice. QFL T cells therefore detect changes in the antigen processing pathway during MCMV infection and serve as an unconventional immune response against infection.

## Results

### MCMV downregulates ERAAP, leading to the presentation of FL9-Qa-1b

Herpesviruses target various proteins in the antigen processing pathway to evade immune responses, including downregulation of ERAP1 mRNA by HCMV (Kim et al., 2011; Romania et al., 2017). To test whether MCMV also targets ERAAP, we measured ERAAP expression in infected cells. We infected B6 fibroblast cells with a GFP expressing MCMV strain for 36 hours and sorted infected (GFP positive cells) and uninfected (GFP negative cells) by fluorescence-activated cell sorting (FACS) (Figure 1A). Western blot analysis in each sorted sample revealed decreased ERAAP protein levels of up to 90% in GFP positive infected cells compared to GFP negative uninfected and mock infected cells (Figure 1B and C). Down-regulation of ERAAP protein was also observed upon MCMV infection of additional cell lines, including RAW macrophages and L cells (Supplemental Figure 1 and 2), indicating the generality of the phenomenon.

**Figure 1:**
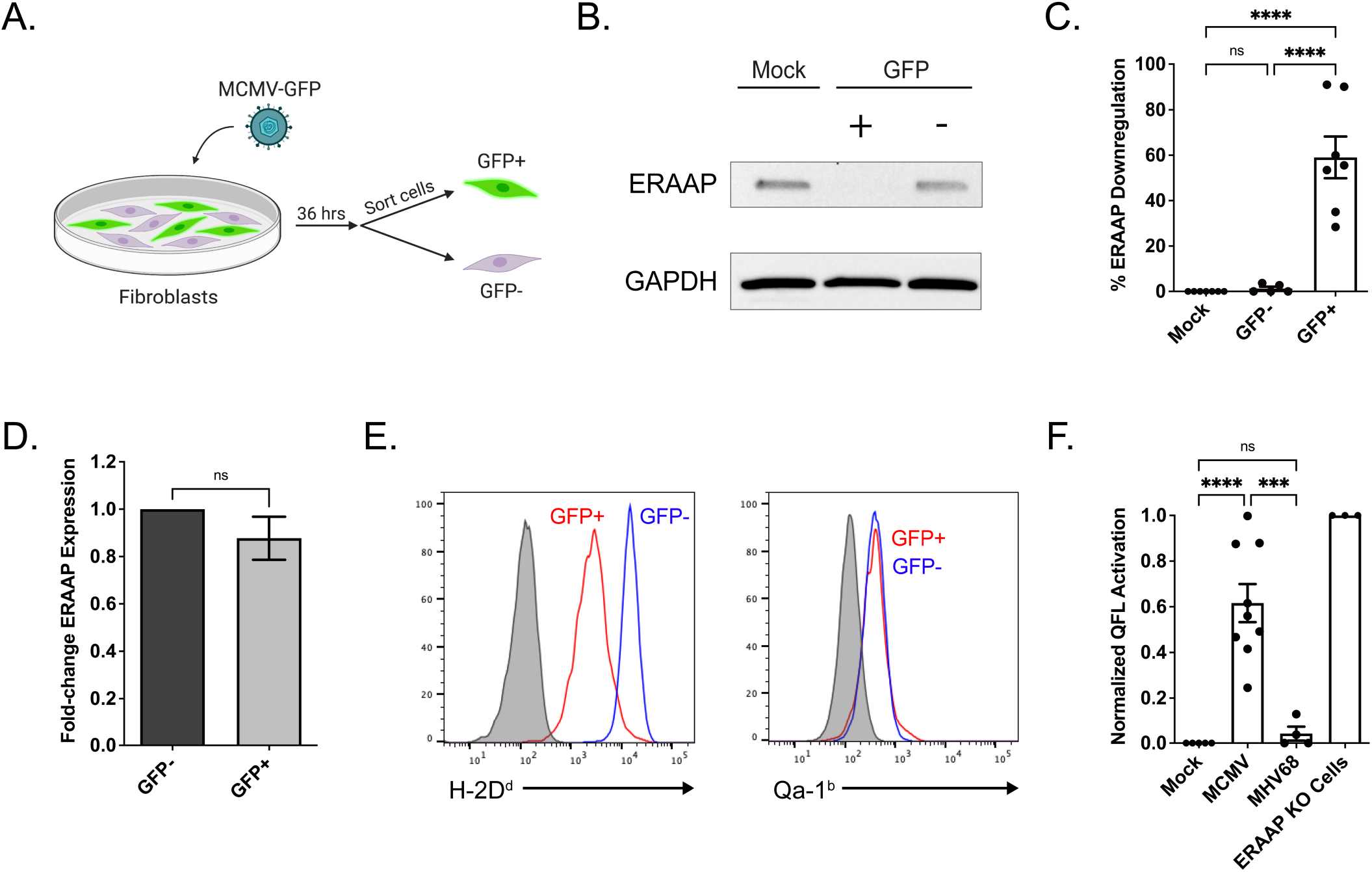
MCMV downregulates ERAAP protein and lead to presentation of the QFL ligand, FL9-Qa1b. (A-D) B6 fibroblast cells were infected with MCMV-GFP and after 36 hours GFP+ and GFP- cells were sorted and assayed by western blot or RT-qPCR. (B) Representative western blot showing ERAAP protein levels in mock infected, GFP positive (+), and GFP negative (-) samples, with GAPDH as the loading control. (C) Compilation of percent ERAAP downregulation in GFP+ and GFP-samples compared to mock infected cells. ERAAP protein band intensity levels were calculated and normalized to the corresponding GAPDH band intensity level, and percent ERAAP downregulation was calculated by comparing normalized ERAAP protein levels in each sample to the mock infected sample. Data shown as mean±SEM. (D) Quantitative real-time PCR analysis of relative ERAAP mRNA levels in GFP+ compared to GFP- sorted cells after normalization to GAPDH. (E) A macrophage cell line (RAW 264.7) was infected with MCMV and analyzed by flow cytometry for surface expression of classical MHC H-2Dd (left panel) and non-classical MHC Qa-1b (right panel) in gated GFP- and GFP+ cells. Experiment is representative of two replicates. Gray histograms represent isotype antibody control. (F) Presentation of the QFL ligand, FL9-Qa1b was measuring the lacZ response of QFL-reactive BEko8Z T cell hybridoma cells incubated with mock infected, MCMV infected, MHV68 infected cells, and ERAAP KO fibroblast cells. The hybridoma response was measured at an OD595 and then normalized to mock infected samples. The percent QFL activation was calculated by comparing the fold-change of the normalized response of each sample to the response against ERAAP knockout cells. Data shown as mean±SEM. Comparisons between samples were made using an ordinary one-way ANOVA multiple comparisons test, *p<0.0332, **p<0.0021, ***p<0.0002, ****p<0.0001.

HMCV infection leads to downregulation of ERAP1 mRNA through viral microRNAs miR-US4-1 and miR-UL112-5p. To determine whether MCMV targeted ERAAP by a similar mechanism, we used RT-qPCR to measure and compare ERAAP mRNA levels in FACS sorted MCMV infected GFP positive and uninfected GFP negative cells. We found that ERAAP mRNA levels in GFP positive infected cells did not change compared to GFP negative uninfected cells (Figure 1D). These data suggest that unlike with HCMV infection, MCMV does not lead to reduced ERAAP mRNA levels and instead inhibits ERAAP at the translational or protein level.

MCMV infection leads to strong MHC Ia downregulation, but whether Qa-1b levels are impacted is not known. To address this question, we infected RAW macrophages with MCMV-GFP and assessed the surface levels of both classical H-2D^d^ (as a representative of classical MHC Ia molecules) and non-classical Qa-1b levels by flow cytometry in GFP negative and GFP positive cells. Classical H-2D^d^ levels were strongly downregulated in infected GFP positive cells, while non-classical Qa-1b levels remained the same compared to GFP negative cells (Figure 1E). We also tested other cells lines such as L cells and found that Qa-1b levels remained unchanged in infected GFP positive versus GFP negative uninfected cells (Supplemental Figure 3). This indicates that MCMV does not alter Qa-1b expression.

Due to the strong downregulation of ERAAP and normal Qa-1b levels observed in MCMV infected cells, we hypothesized that these cells would present a peptide repertoire similar to one observed with ERAAP knockout cells. Thus, we investigated whether infected cells presented the QFL ligand (FL9 peptide bound to Qa-1b). To do so, we used an FL9-Qa-1b reactive T cell hybridoma cell line (BEko8Z) with a TCR inducible LacZ gene (Nagarajan et al., 2012) and L cells expressing Qa-1b as antigen presenting cells, and tested the ability of MCMV infected L cells to activate the T cell hybridomas (Figure 1F). L cells were infected with MCMV for 36 hours before incubation with BEko8Z cells for 24 hours. We determined BEko8Z activation, as indicated by the induction of LacZ, by measuring colorimetric cleavage of the beta-galactosidase substrate CPRG. BEko8Z cells responded strongly to MCMV infected L cells and did not respond to L cells infected with a different herpesvirus MHV68. These results indicate that MCMV infection leads to presentation of the FL9 peptide on Qa-1b, likely due to viral downregulation of ERAAP protein levels.

### MCMV infection induces CD8^+^ T cell responses that target ERAAP KO cells

If ERAAP downregulation occurred *in vivo* during MCMV infection, we might expect MCMV infected mice to generate an effector CD8^+^ T cell response against ERAAP KO cells, similar to that observed upon immunization of mice with ERAAP ko splenocytes (Hammer et al., 2007; Nagarajan et al., 2012). To examine this question, we infected mice with MCMV, and 10 days later, isolated splenocytes from infected mice and re-stimulated them *ex vivo* with WT or ERAAP KO spleen cells (Figure 2A). We intracellularly stained and analyzed samples by flow cytometry to measure levels of IFN-γ^+^ CD8^+^ T cells. Our experiments indicate that CD8^+^ T cells from MCMV infected mice were indeed capable of mounting a strong IFN-γ response against ERAAP KO cells. About 4% of CD8^+^ T cells in MCMV infected mice induced IFN-γ against ERAAP KO cells, while no IFN-γ response was seen against WT cells (Figure 2B and C). This was comparable to the response of CD8^+^ T cells from WT mice immunized with ERAAP KO spleen cells (ERAAP KO immunized), where 7% of CD8^+^ T cells induced IFN-γ^+^ against ERAAP KO cells. No IFN-γ^+^ response was seen in control WT mice immunized with WT spleen cells (uninfected).

**Figure 2:**
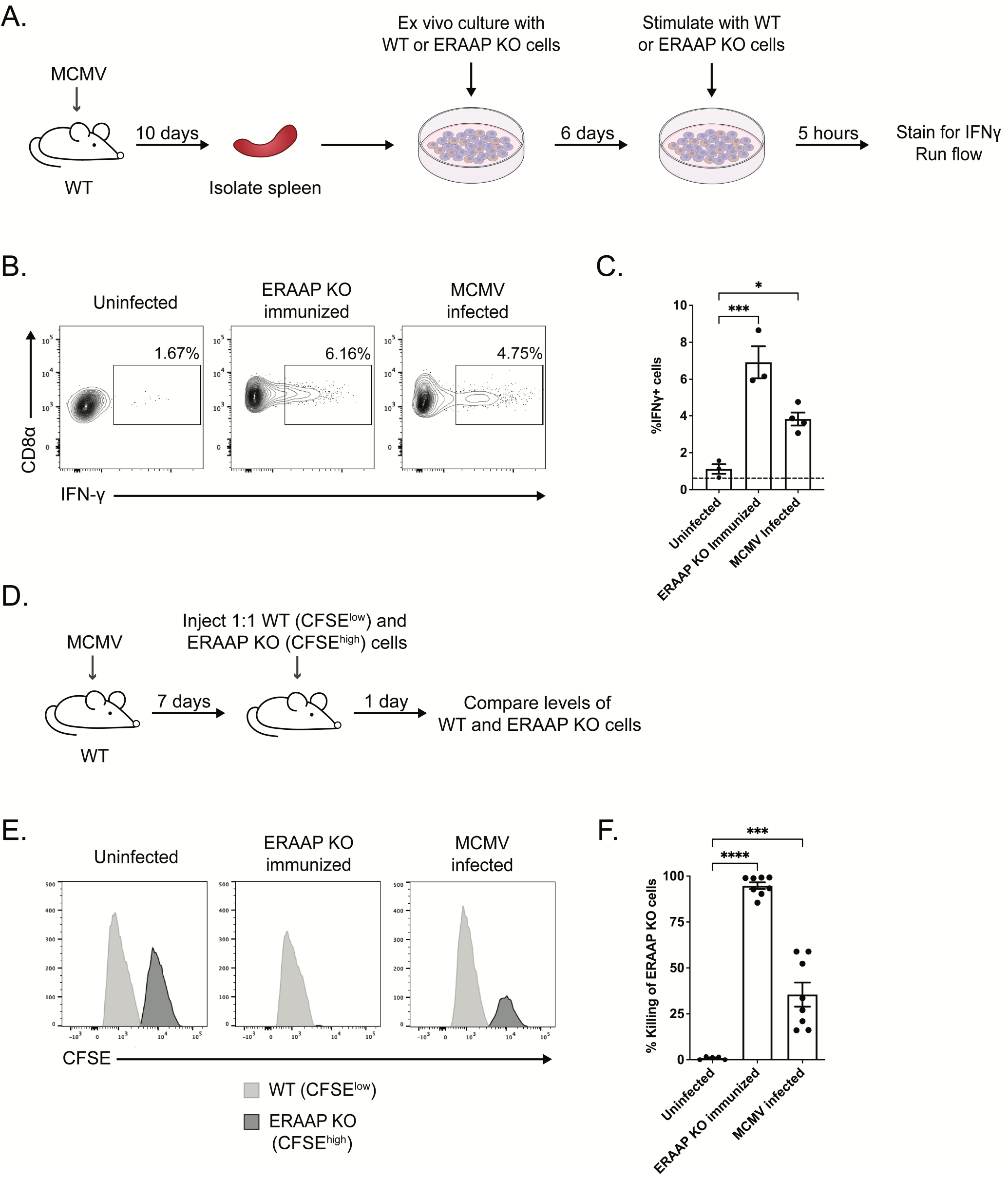
MCMV infection stimulates an response to ERAAP deficient cells. (A-C) C57Bl/6 (WT) mice were infected with MCMV (MCMV infected), immunized with ERAAP knockout splenocytes (ERAAP KO immunized), or immunized with WT splenocytes (uninfected). At 10 days post infection, splenocytes were isolated from mice and restimulated *ex vivo* with WT or ERAAP KO splenocytes for 6 days. Cells were stimulated with WT or ERAAP KO splenocytes for 5 hours to measure intracellular IFN-γ levels. (B) Representative flow cytometry plots of IFN-γ producing CD8α+ cells from uninfected, ERAAP KO immunized, and MCMV infected mice against ERAAP KO target spleen cells. (C) Compiled data of %CD8α+IFN-γ+ cells in each group stimulated against ERAAP KO cells. Dotted line represents average %CD8α+IFN-γ+ cells against WT target spleen cells. Data shown as mean±SEM. (D-F) C57Bl/6 (WT) mice were infected with MCMV. At 7 days post infection, infected mice were injected with splenocytes from WT or ERAAP KO mice that had been CFSE labelled with 0.5 μM (CFSE^low^) and 5 μM (CFSE^high^), respectively., ERAAP KO immunized, and WT immunized (uninfected) mice. (E) Representative flow cytometry plots showing killing of CFSE^high^ ERAAP KO targets in each group. (F) Compiled data for percent killing of ERAAP KO target cells. Percent killing was calculated as follows: 1-(%CFSE^hi^ sample / %CFSE^lo^ sample) / (%CFSE^hi^ uninfected / %CFSE^lo^ uninfected) x 100. Data shown as mean±SEM. Comparisons between samples were made using an ordinary one-way ANOVA multiple comparisons test, *p<0.0332, **p<0.0021, ***p<0.0002, ****p<0.0001.

We then assessed the ability of the immune response elicited in MCMV infected mice to target and specifically lyse ERAAP KO spleen cells using an *in vivo* killing assay (Figure 2D). WT mice were infected with MCMV for 7 days and then injected with a 1:1 mixture of splenocytes from WT and ERAAP KO mice labeled with two different concentrations of proliferation dye CFSE. After 24 hours, the proportion of WT versus ERAAP KO populations were compared to measure percent killing in infected mice. Labeled target cells were also injected into uninfected mice immunized with WT cells or immunized with ERAAP KO cells (ERAAP KO immunized) as negative and positive controls, respectively. As predicted, MCMV infection induced a cytotoxic T cell response that efficiently targeted up to 60% of ERAAP KO target cells (Figure 2E and 2F). ERAAP KO immunized mice had a more robust response, resulting in an average of 94% killing of ERAAP KO cells. We conclude that ERAAP downregulation during MCMV infection elicits a potent immune response that specifically induces IFN-γ against and eliminates ERAAP KO cells *in vivo*.

### QFL T cells proliferate in response to MCMV infection

QFL T cells are found at a relatively high frequency and exhibit a mostly CD44+ phenotype in wild type mice, but expand further and acquire effector activity upon immunization with ERAAP deficient splenocytes (Nagarajan et al., 2012). We investigated whether QFL T cells expanded during MCMV infection *in vivo* in response to ERAAP downregulation by the virus. We infected mice with MCMV for 10 days and determined the number of splenic QFL T cells using tetramer enrichment and flow cytometry (Figure 3A). Uninfected mice immunized with WT cells or immunized with ERAAP KO cells (ERAAP KO immunized) served as negative and positive controls, respectively. Our results revealed that MCMV infected mice had a three-fold increase in the total number of QFL T cells in the spleen compared to uninfected mice. Specifically, MCMV infected mice had an average of 3100 total QFL T cells in the spleen, while uninfected mice had an average of 1000 total QFL T cells in the spleen, consistent with previous studies (Figure 3B and 3C) (Nagarajan et al., 2012). In addition, all tetramer^+^ QFL T cells expressed the Vα3.2 chain as previously shown (Guan et al., 2017).

**Figure 3:**
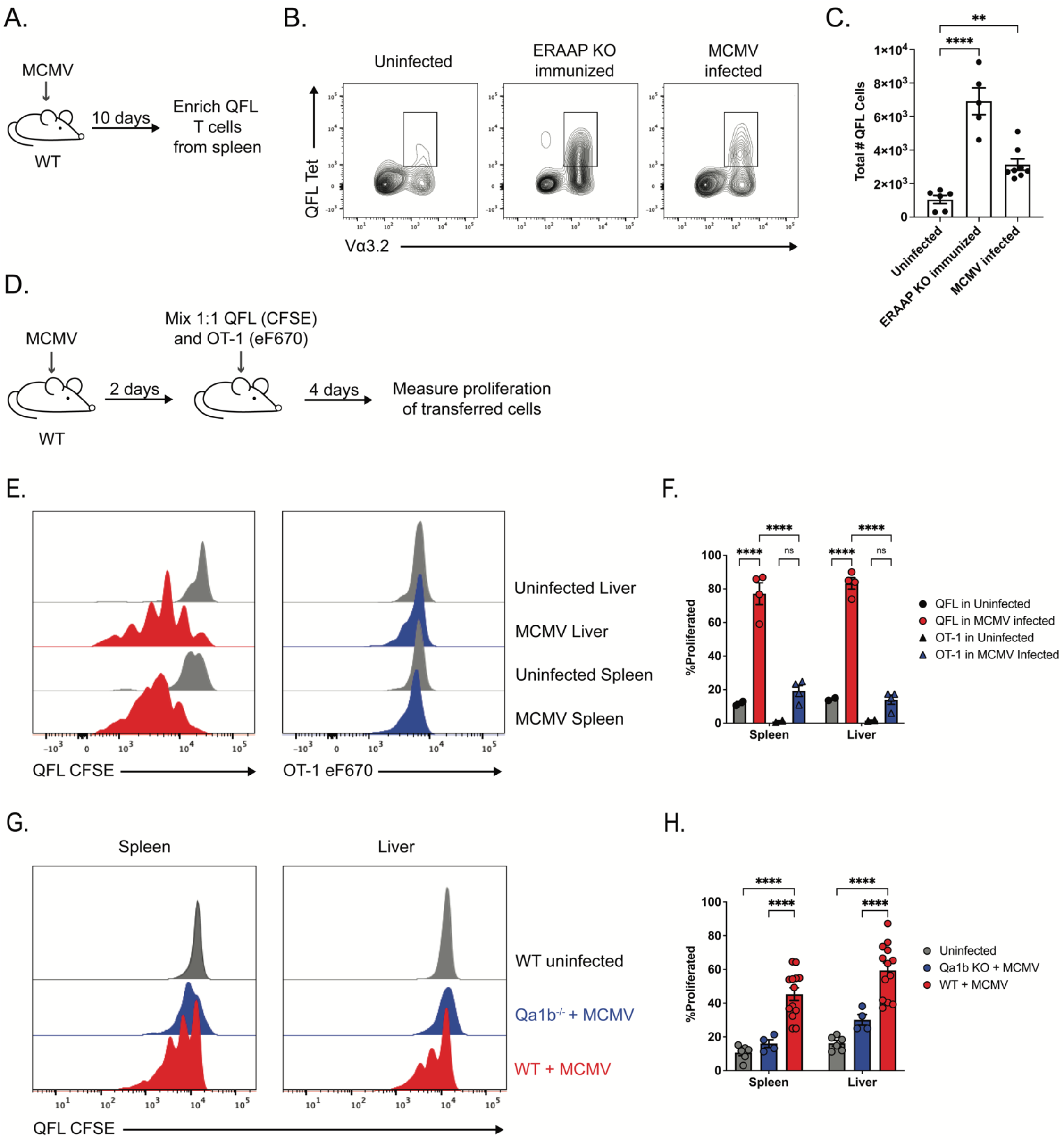
QFL T cells expand and proliferate during MCMV infection. (A-C) WT mice were uninfected, immunized with ERAAP knockout cells (ERAAP KO immunized), or infected with MCMV (MCMV infected) and QFL T cells were stained with FL9-Qa-1b tetramers and enriched from the spleen 10 days post infection. (B) Representative flow cytometry plots showing QFL expansion by QFL tetramer and Va3.2 expression in each group. (C) Compiled data of total number of QFL T cells enriched from the spleen in each group. Data shown as mean±SEM. (D-F) WT mice were uninfected or MCMV infected. Two days post infection, QFL cells (CFSE) and OT-1 cells (eF670) were labelled, mixed 1:1, and transferred intravenously into recipient mice. Four days post transfer, proliferation of both T cell subsets was measured in the liver and spleen. (E) Representative flow cytometry plots of QFL (left panel) and OT-1 (right panel) cell proliferation in the spleen and liver of uninfected and MCMV infected mice. (F) Compiled data of the percentage of proliferated QFL and OT-1 cells in the spleen and liver of uninfected and MCMV infected mice. Data shown as mean±SEM. (G) Representative flow cytometry plots of levels of proliferation of transferred QFL T cells into WT uninfected, Qa1b knockout MCMV infected, and WT MCMV infected mice. WT and Qa-1b knockout mice were infected with MCMV. Two days post infection, QFL cells (CFSE) were labelled and transferred intravenously into recipient mice. Four days post transfer, proliferation was measured in the spleen (left panel) and liver (right panel). (H) Compiled data of the percentage of proliferated QFL cells in the spleen and liver of WT uninfected, Qa1b knockout MCMV infected, and WT MCMV infected mice. Data shown as mean±SEM. Comparisons between samples were made using an ordinary two-way ANOVA multiple comparisons test, *p<0.0332, **p<0.0021, ***p<0.0002, ****p<0.0001.

To measure in vivo presentation of the QFL ligand (FL9-Qa-1b), we generated a transgenic mouse line harboring the rearranged TCR alpha and beta genes from the FL9-Qa-1b reactive Beko8Z hybridoma (QFL TCR transgenic mice). We transferred CFSE labeled splenocytes from QFL TCR transgenic mice into WT mice that had been infected 2 days prior with MCMV and measured the proliferation of transferred QFL T cells in different tissues 4 days after transfer (Figure 3D). We focused on the liver and spleen of infected mice, which are known sites of acute MCMV infection (Hsu et al., 2009). To confirm that QFL proliferation was due specifically to FL9-Qa-1b presentation and not due to a non-specific inflammatory response to viral infection, we also co-transferred eF670 labeled transgenic OT-1 T cells, specific for the OVA-K^b^ ligand which is not present in our system (Hogquist et al., 1994; Shinkai et al., 1992). We found that in MCMV infected mice, an average of 77% of transferred QFL T cells proliferated in the spleen and 83% of QFL T cells proliferated in the liver (Figure 3E and F). OT-1 T cells proliferated at significantly lower levels, where about 19% and 13% of transferred OT-1 cells proliferated in the spleen and liver, respectively. In addition, QFL T cell proliferation induced by MCMV infection was significantly reduced in Qa-1b deficient (Qa-1b^-/-^) MCMV infected mice (Figure 3G and H), further confirming the specificity of this response. These data suggest that QFL T cells encounter their high affinity ligand in the liver and spleen of MCMV infected mice, leading to their proliferation and expansion.

### QFL T cells acquire an effector phenotype upon MCMV infection

We next investigated how MCMV infection impacted the activation phenotype of QFL T cells. To characterize the effector phenotype and potential anti-viral role of these QFL T cells during MCMV infection, we examined expression of KLRG1, an effector marker expressed on cytotoxic, proliferative, and virus-specific T cells (Herndler-Brandstetter et al., 2018; McMahon et al., 2002; Thimme et al., 2005), along with CD44: a marker of antigen experience, on QFL T cells in mice 10 days after infection with MCMV (Figure 4). As hypothesized, QFL T cells further upregulated CD44 from 53% in uninfected to 89% in MCMV infected mice (Figure 4A and B). QFL T cells also upregulated KLRG1 from 0% in uninfected to 43% in MCMV infected mice (Figure 4C and D). CD44 and KLRG1 levels on QFL T cells from MCMV infected mice were comparable to the levels found in ERAAP KO immunized mice. Thus, QFL T cells expand and differentiate into effector and cytotoxic T cells during MCMV infection.

**Figure 4:**
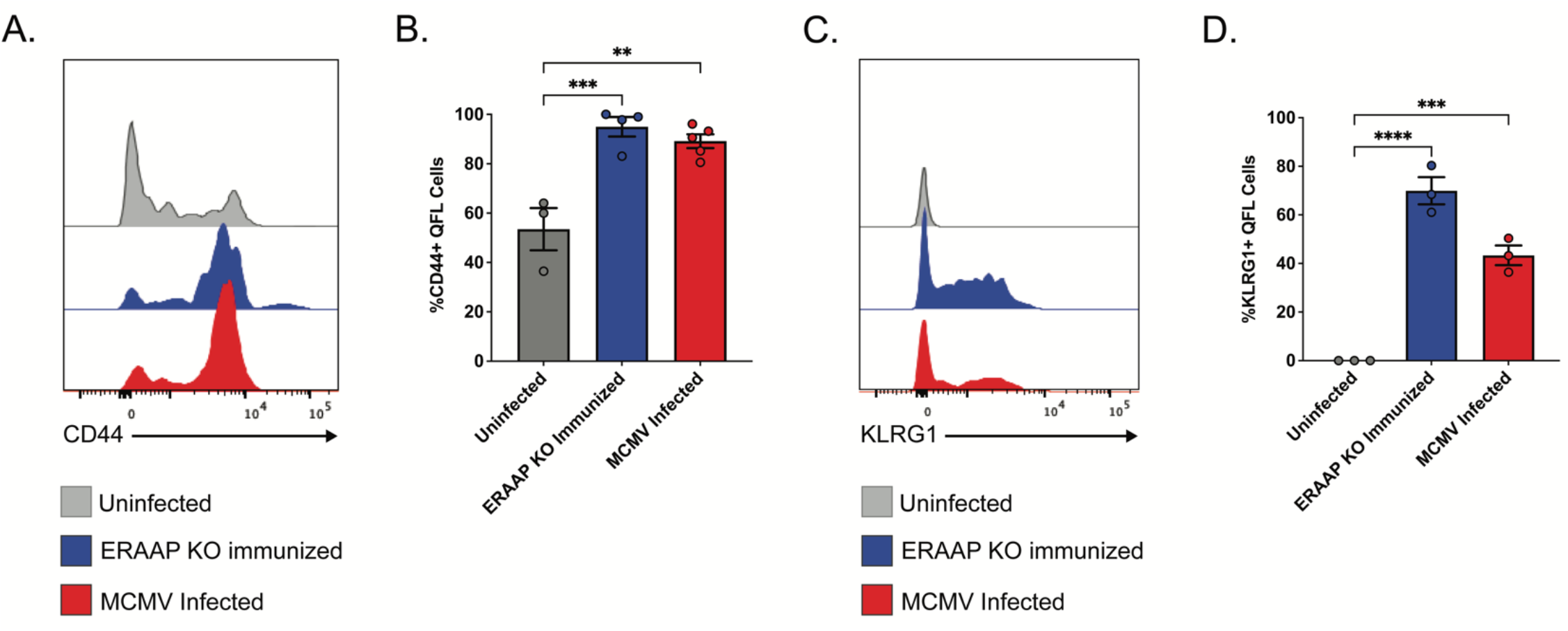
QFL T cells upregulate effector markers CD44 and KLRG1 in MCMV infected mice. (A-B) WT mice were uninfected, ERAAP KO immunized, or MCMV infected. QFL T cells were enriched from the spleen of each mouse and CD44 and KLRG1 expression was analyzed. (A) Representative flow cytometry plots for CD44 expression in QFL T cells. (B) Compiled data of %CD44+ QFL T cells for each group. Data shown as mean±SEM. (C) Representative flow cytometry plots for KLRG1 expression in QFL T cells. (D) Compiled data of %KLRG1+ QFL T cells for each group. Data shown as mean±SEM. Comparisons between groups were made using an ordinary one-way ANOVA multiple comparisons test, *p<0.0332, **p<0.0021, ***p<0.0002, ****p<0.0001.

### QFL T cells provide protection against severe MCMV infection

Due to the expansion and effector differentiation of QFL T cells during MCMV infection, we predicted that QFL T cells could provide protection against MCMV infection. To test this, we adoptively transferred transgenic QFL T cells into immunodeficient Rag2/γc knockout mice, which lack functional T, B, and NK cells, and infected the mice with MCMV 2 days post transfer. The viral load was assessed 12 days after infection (Figure 5A). As a control, we transferred transgenic OT-1 T cells (specific for OVA, which is not present in our system) into Rag2/γc knockout mice to compare the anti-viral response of a non-specific transgenic T cell response to the QFL T cell response. qPCR analysis of the MCMV titers in the spleen and liver of infected QFL recipient mice showed significant reduction of viral load in both tissues. Immunodeficient mice that received QFL T cells had 2-log reduction in MCMV titers in the spleen (Figure 5B) and liver (Figure 5C) compared to the OT-1 recipient mice. Therefore, QFL T cells provided effective control of MCMV in Rag2/γc knockout mice.

**Figure 5:**
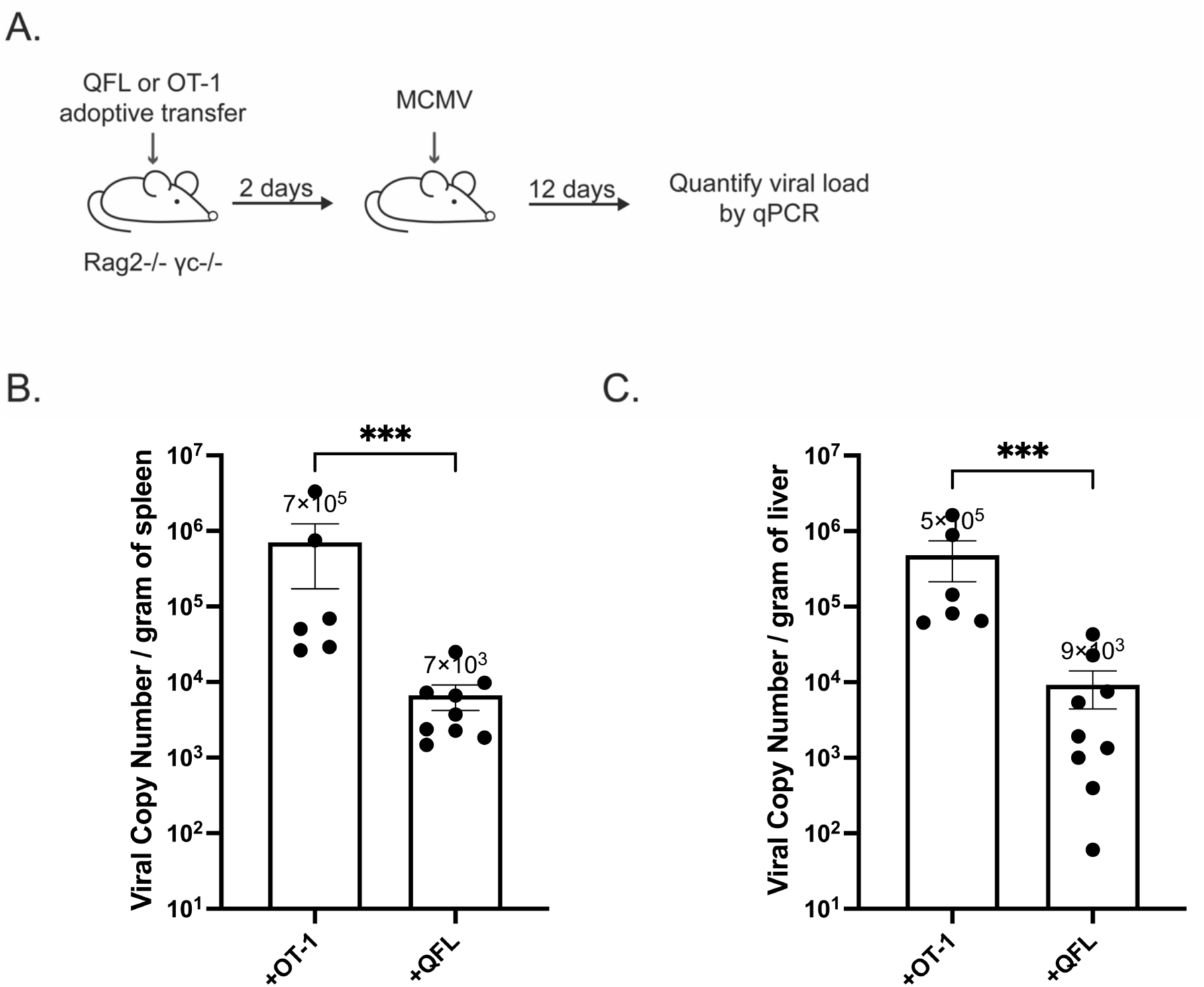
QFL T cells provide protection against MCMV infection. (A-C) 2.5e5 QFL or OT-1 T cells from TCR transgenic mice were adoptively transferred into Rag2/γc knockout mice. After 2 days post transfer, mice were infected with MCMV. After 12 days post infection, the liver and spleen were isolated from infected mice and homogenized. MCMV titers were determined by qPCR (B) Compiled data of viral titers in the spleen of infected Rag2/γc knockout mice adoptively transferred with either QFL or OT-1 cells. Data shown as mean±SEM. (C) Compiled data of viral titers in the liver of infected Rag2/γc knockout mice adoptively transferred with either QFL or OT-1 cells. Data shown as mean±SEM. Comparisons between samples were made using Mann-Whitney test, ***p<0.0002.

## Discussion

Viruses and their hosts have co-evolved for millions of years, shaping viral immune evasion strategies and compensatory immune responses from the host. For example, CMV downregulates MHC Ia molecules to avoid CTL attack, leading to enhanced recognition by NK cells. Here we show that MCMV also triggers downregulation of the MHC I peptide processing enzyme ERAAP in infected cells, and that mice infected with ERAAP develop a unconventional CTL response directed against uninfected ERAAP deficient cells. Our data fit with a model (Supp Fig 4) in which MCMV targets ERAAP in order to evade conventional CTL responses. Given that conventional anti-viral T cells to MCMV are primarily primed through cross-presentation of viral peptides by uninfected dendritic cells with normal levels ERAAP (Busche et al., 2013; Cruz et al., 2017), downregulation of ERAAP in infected cells would contribute to evasion of conventional T cells by altering the peptide display in cells harboring virus. However, the immune system counters by arming unconventional CTL to eliminate cells with dysfunctional ERAAP. Thus both viral downmodulation of ERAAP and the immune response to loss of ERAAP are part of the layered interplay between MCMV and the host immune system.

Previous studies identified a unconventional T cell response to a self-peptide presented by Qa1b (QFL T cells) that makes up a prominent part of the T cell response to ERAAP deficient cells in wild type mice (Nagarajan et al., 2012). Here we show that MCMV infection triggers the expansion and effector differentiation of QFL T cells, raising the question of which cells serve as APC for QFL T cells in infected mice. Given that the majority of QFL T cells in uninfected mice have an antigen-experienced (CD44+) phenotype, they may not require priming by an activated DC, as do naïve T cells. In addition, since Qa1b and Fam49A/B are broadly expressed, QFL T cells could potentially expand and differentiate upon encounter with any virally infected cell. For example, in liver, where we observed substantial QFL T cell proliferation in infected mice, virus can replicate in liver sinusoidal cells, including endothelial cells and Kupffer cells (Kern et al., 2010; Salazar-Mather et al., 1998; Tegtmeyer et al., 2019). In addition to activating QFL T cells, virally infected cells may also serve as targets for killing, helping to explain the protective effect of QFL T cells on viral control reported here.

Protective Qa-1b restricted CD8^+^ T cell responses have been described in several infection models including *Mycobacterium tuberculosis, Listeria monocytogenes*, and *Salmonella typhimurium*, and MCMV (Anderson et al., 2019; Bian et al., 2017; Lo et al., 1999; Seaman et al., 1999; Shang et al., 2016). While thus far these responses appeared to be directed against microbial peptides, we show here that a response to self-peptides can form an important part of these protective responses. In that regard, down regulation of ERAAP could serve as a kind of “danger signal” alerting the host to disruption of cellular homeostasis by altering peptide display on MHC-E. T cells reactive to this form of altered self may not be overtly self-reactive, due to the predominant presentation of Qdm peptides and low level of expression of MHC-E on normal cells. However, they may nevertheless acquire a pre-activated/memory phenotype due to low level or infrequent encounters with altered self on stressed or abnormal cells, and thus may respond more rapidly upon infection. It is tempting to speculate that such a preformed T cell response may enhance other anti-viral non-classical T cell responses, including those directed against viral peptides. Of particular interest is the broad and protective CD8^+^ response restricted to MHC-E elicited by a rhesus macaque CMV vectored SIV vaccine (Hansen et al., 2011, 2013a, 2013b, 2016, 2019; Malouli et al., 2021; Verweij et al., 2021; Yang et al., 2021). While it is not yet known if RhCMV downregulates ERAAP, it is possible that a QFL-like response to self may contribute to the potent anti-viral MHC-E restricted response in this setting.

HCMV is ubiquitous worldwide, where 80-90% of the global population is seropositive, and the virus is the leading infectious cause of congenital neurological diseases and deafness (Crough and Khanna, 2009; Elemans et al., 2012; Heiden et al., 2014). Our work is applicable to HCMV models and suggests there could be significant value in identifying an HLA-E-restricted QFL-like T cell response that detects ERAAP downregulation and contributes to protection against HCMV infection. Our findings can also be applied to other infectious diseases, such as tuberculosis and HIV, which also induce CTL responses restricted to non-classical MHC. In addition, several cancer models have also been shown to have attenuated ERAAP levels (Kamphausen et al., 2010), leading to presentation of immunogenic tumor antigens and enhanced antitumor immunity (James et al., 2013). The role of QFL T cells in different infectious disease and anti-tumor models remains to be determined but seems to be a promising path to explore. Specifically, a vaccine that was designed to be directed towards a specific non-classical MHC, such as FL9-Qa-1b, could be used universally to stimulate similar CTL responses across individuals, despite potential patient-to-patient differences in MHC I genotype.

## Acknowledgements

This work is dedicated to Nilabh Shastri, who was a pioneer in immunology, antigen presentation and processing, and an amazing mentor and friend. We thank H. Nolla and A. Valero of the UC Berkeley Cancer Research Lab for help with flow cytometry. We would like to thank the Shastri lab, Robey lab, and Glaunsinger lab for helpful and creative discussions about this project. Supplemental figure 4 was created with BioRender.com. Funding was provided by National Institutes of Health (RO1AI149341). K.M.G and M.M. were supported by National Science Foundation Graduate Research Fellowships.

## Author contributions

K.M.G., M.M., N.S., E.R., and L.C. conceived and designed the experiments. K.M.G., R.C., and C.P. performed the experiments. K.M.G. performed data analyses. N.S., E.R., and L.C. provided funding, reagents, materials, and analysis tools. A.Y.L generated the transgenic mouse strain. K.M.G., M.M., R.C., C.P., E.R., and L.C. contributed to writing this paper.

---

**Supplemental Figure 1: ERAAP downregulation in MCMV infected RAW macrophages** Representative western blot showing ERAAP protein levels in GFP positive (+) and GFP negative (-) samples, with GAPDH as the loading control. RAW macrophages were infected with MCMV-GFP and after 36 hours GFP+ and GFP-cells were sorted and assayed by western blot.

**Supplemental Figure 2: ERAAP downregulation in MCMV infected L cells**

Representative western blot showing ERAAP protein levels in GFP positive (+) and GFP negative (-) samples, with GAPDH as the loading control. L cells were infected with MCMV-GFP and after 36 hours GFP+ and GFP-cells were sorted and assayed by western blot.

**Supplemental Figure 3: Qa-1b levels remain unchanged in MCMV infected L cells**

Representative flow cytometry plots showing surface expression of non-classical MHC Qa-1b in GFP- (blue) and GFP+ (red) MCMV infected L cells. Gray histograms represent isotype antibody control.

**Supplemental Figure 4: Model for ERAAP downregulation and QFL activation during MCMV infection**

During MCMV infection, cross-presenting dendritic cells (DCs) with functional ERAAP and MHC I prime conventional CD8^+^ T cell responses against MCMV. These primed T cells, however, cannot identify and target MCMV infected cells since the infected cells have downregulated ERAAP and present an altered peptide repertoire. Viral peptides may also not get presented due to ERAAP downregulation, and ones that are presented may have a lower affinity. As a result, MCMV evades conventional T cell responses, and lowers the efficiency of responses that do recognize infected cells. However, by downregulating ERAAP, this causes the presentation of FL9 on Qa-1b, thus inducing QFL T cells to expand in response to this ligand. As a result, these QFL T cells recognize and target infected cells and play an anti-viral role during MCMV infection.

## Methods

### Mice, immunizations, and infections

WT C57Bl/6 mice were purchased from the Jackson Laboratory. ERAAP-KO and H2-T23 Qa-1b KO mice have been described (Hammer et al., 2006; Nagarajan et al., 2016). Rag2/OT-I transgenic knockout mice were purchased from Taconic. All experiments were done using 7-10 week old mice. Uninfected mice were immunized with 2e7 WT splenocytes, and ERAAP-KO immunized mice were immunized with 2e7 ERAAP-KO splenocytes intraperitoneally. Infected mice were infected with 1e6 PFU of WT MCMV intraperitoneally. All mouse experiments were done with the approval of Institutional Animal Care and Use Committee of the University of California, Berkeley.

### Cell lines

B6 fibroblasts were generated by the Shastri lab from B6 mice. Qa-1b expressing L cells have been described (Nagarajan et al., 2012). NIH 3T3 (ATC#CRL-1658, RRID:CVCL_0594)) and RAW 264.7 mouse macrophages (ATCC TIB-71, RRID:CVCL_0493) were purchased from ATCC. QFL hybridomas (BEko8Z cells) were established in the Shastri lab as described (Nagarajan et al., 2012). B6 fibroblasts and splenocytes were cultured in RPMI (cRPMI) with 10% FBS (Invitrogen, Carlsbad CA), 100 U/ml Penicillin/Streptomycin (Invitrogen), 2mM L-Glutamine, and 50 μM 2-ME. NIH 3T3 and RAW 264.7 cells were cultured in DMEM with 10% FBS (Invitrogen, Carlsbad CA) and 100 U/ml Penicillin/Streptomycin (Invitrogen).

### Generation of QFL transgenic mice

The QFL transgenic mice were generated on the B6 background in the Cancer Research Laboratory Gene Targeting Facility at UC Berkeley under standard procedures. The QFL TCR alpha and beta chain sequences were previously identified and amplified from the genomic DNA of QFL BEko8Z hybridoma cells (Guan et al., 2017; Nagarajan et al., 2012). The TRAV9D-3 TCR alpha chain was cloned with the forward primer (5’ AAAACCCGGGCCAAGGCTCAGCCATGCTCCTGG) with an added XmaI cutting site at the 5’ end of the DNA sequence and a reverse primer for TRAJ21 (5’ AAAAGCGGCCGCATACAACATTGGACAAGGATCCAAGCTAAAGAGAACTC) with an added Not1 cutting site at the 5’ end of the DNA sequence. The TCR beta chain was cloned with the forward primer (5’ AAAACTCGAGCCCGTCTGGAGCCTGATTCCA) with an added Xho1 cutting site at the 5’ end of the DNA and a reverse primer for TRBJ2-7 (5’ AAAACCGCGGGGGACCCAGGAATTTGGGTGGA) with a SacII cutting site flanking the 5’ end of the DNA sequence. The cloned TCR alpha chain was cloned into pTa cassette vector using the Xmal and Not1 restriction sites, while the TCR beta chains were cloned into pTb cassette vector using Xhol and SacII restriction sites (Kouskoff et al., 1995). The ampicillin resistance gene was removed from the pTa and pTb cassettes by digestion with EarI. The QFL mice were maintained on the B6 background and bred once with B6 Ly5.1 mice to generate (QFLTgxB6 Ly5.1/2) background mice for use in experiments. Founder mice were identified by flow cytometry and PCR genotyping of tail genomic DNA using the primers mentioned above.

### QFL Hybridoma BEko8Z Assay

Hybridoma cells were used in a LacZ assay as described (Sanderson and Shastri, 1994). Briefly, fibroblasts expressing Qa-1b were plated at 20,000 cells/well in a 96-well plate and infected with MCMV at an MOI 10 for 24 hours before addition of 1e5 QFL hybridoma cells. After 16 hours, cells were spun down at 1500rpm for 5 minutes before addition of a fluorescent β-galactosidase substrate. The absorbance of each well was examined 12 hours after incubation at a wavelength of 595nm.

### Virus production and propagation

E. coli strain GS1783 containing MCMV BAC pSM3fr was provided by Dr. Caroline Kulesza (Fort Lewis College). NIH 3T3 cells were used to produce and titer virus by TCID50. Mutant MCMV lacking ORFs m1-22 (MCMV Δm1-22) was a gift from Dr. Hidde Ploegh (Whitehead Institute, MIT, MA). MCMV GFP virus was generously provided by Dr. Koszinowski (Max von Pettenkofer-Institute, Munich, Germany).

### Western blot

B6 fibroblasts were infected with MCMV-GFP for 36 hours and then sorted by FACS. Mock infected and GFP+/GFP- sorted cells were lysed in protein lysis buffer 25 mM Tris-HCl (pH 7.6), 150 mM NaCl, 0.1% SDS, 0.5% sodium deoxycholic, and complete EDTA-free protease inhibitors (Roche) for 30 minutes on ice. After, samples were centrifuged at 20,000xg for 10 min at 4°C to clarify lysate. Lysates were separated by SDS-PAGE and western blotted with rabbit anti-ERAAP (1:1000) or mouse anti-GAPDH (1:5000; Abcam, ab8245). Rabbit anti-ERAAP was produced in the Shastri lab.

### RT-qPCR and qPCR

qPCR was conducted using an ABI7300 RT-qPCR System with the following protocol: 95°C dissociation step for 15 sec, 60°C amplification step for 1 min, repeated for 40 cycles. The Applied Biosystems 7300 SDS software was used to calculate Cq values. Each sample was done in triplicate and averaged.

DNA and RNA extraction were done as previously described (Greene et al., 2016). Mouse and viral DNA were isolated from mouse tissue using the Qiagen DNeasy Blood and Tissue Kit (Qiagen). iTAQ universal Syber Green supermix (Invitrogen) and 300uM of MCMV Gb or GAPDH primers were mixed with isolated DNA for qPCR analysis. Absolute viral copy numbers were extrapolated using a standard curve of known quantities of purified MCMV BAC.

RNA was extracted from cells using Trizol (Invitrogen). Genomic DNA was removed with 0.002 U of DNase I (Thermo). cDNA was synthesized using 1 ug of RNA reverse transcribed for 50 min at 42°C using oligo(dT) primer (IDT) and SuperScript II RT (Invitrogen). For qPCR analysis, 2 ul of prepared cDNA was mixed with iTAQ universal Syber Green supermix (Invitrogen) and 300uM of ERAAP primers. ERAAP mRNA levels were compared by ΔΔCT using average Cq values normalized to GAPDH.

### Spleen and liver cell isolation

Spleen cells from mice were homogenized in PBS using a GentleMACS Dissociator (Miltenyi Biotec), washed in FACS buffer (PBS+0.1%sodium azide+5% FCS), and passed through a 70um filter. Cells were RBC lysed with ACK lysis buffer (0.15M NH_4_CL, 1mM KHCO_3_, 0.1mM Na_2_EDTA). Livers were perfused with PBS and homogenized in PBS using a GentleMACS Tissue Dissociator (Miltenyi Biotec). Liver samples were then incubated with 0.1% collagenase type 1A and 40 ug/mL of DNase I for 20 minutes at 37°C before washing with cRPMI and passing through a 70uM filter. Lymphocytes were isolated using a Percoll gradient of 40% percoll layer overlaid onto a 70% percoll layer after spinning at 500xg for 30 minutes at room temperature.

### Antibodies and flow-cytometry

Antibody staining was done in 2.4G2 media with cell surface antibodies for 30 minutes on ice. The following antibodies and clones were used for staining: CD8a Clone 53-6.7, B220 Clone RA3-6B2, Va3.2 Clone RR3-16, KLRG1 Clone 2F1/KLRG1, CD44 Clone IM7, CD8b Clone YTS156.7.7, CD45.1 Clone A20, and IFNy Clone XMG1.2.

### *Ex vivo* T cell restimulation assay

Spleens were harvested from mice 10 days post immunization or infection, and 5e6 spleen cells were restimulated *in vitro* with 5e6 irradiated ERAAP KO spleen cells and 20 U/ml of recombinant human IL-2 (BD Biosciences). After 6 days, cells were isolated and were stimulated for 5 hours with CD4 and CD8 depleted WT or ERAAP KO spleen cells. Golgi-Plug (BD Biosciences) was added after 4 hours of stimulation. Cells were stained with surface markers, fixed, permeabilized (Cytofix/Cytoperm Kit BD Biosciences), and stained intracellularly for IFN-γ.

### *In vivo* cytotoxicity assay

WT mice were infected with 1e6 PFU of WT MCMV intraperitoneally. For controls, mice were immunized with 2e7 of WT or ERAAP KO spleen cells intraperitoneally. WT and ERAAP-KO spleen cells were isolated and stained with 0.5uM or 5uM of CFSE dye (ThermoFisher), respectively. Cells were counted and mixed at a 1:1 ratio and injected intravenously into immunized and infected mice. After 24 hours, spleen cells from each mouse were isolated and the ratio of CFSE^lo^ (WT cells) to CFSE^hi^ (ERAAP-KO cells) was measured by flow cytometry. Percent killing was calculated as follows: 1-(%CFSE^hi^ sample / %CFSE^lo^ sample) / (%CFSE^hi^ uninfected / %CFSE^lo^ uninfected) x 100.

### *In vivo* antigen presentation assay

WT mice were infected with 1e6 PFU of MCMV intraperitoneally. Two days later, QFL and OT-1 transgenic T cells were labeled with 5uM CFSE or 5uM eF670, respectively. Labeled cells were mixed 1:1 and 2e6 cells were injected into infected mice. After 4 days, the liver and spleen were isolated and proliferation was analyzed by flow cytometry.

### QFL T cell Enrichment

QFL T cells were enriched using a QFL tetramer (FL9-Qa-1b) synthesized by the NIH tetramer core facility. Spleen cells were isolated as stated above and incubated with 50 nM of Dasatinib (Cell signaling #9052) at 37°C for 30 minutes. Cells were then washed and incubated with PE-QFL tetramer (1:200) for 1 hour at RT. Cells were washed, resuspended in 150ul FACS buffer with 100 ul of anti-PE microbeads (Miltenyi Biotec), and incubated for 20 minutes at 4°C. Cells were then washed and passed through an LS magnetic column (Miltenyi Biotec). Enriched cells were stained with antibodies for B220, CD8α, CD8β, and Vα3.2. Tetramer+ QFL T cells were gated as B220^-^ CD8α^+^QFL Tet^+^Vα3.2^+^. CountBright beads (Invitrogen) were used in each sample to measure total cell numbers in the enriched and unenriched samples.

### Statistical Analysis

Statistical analysis for each experiment was done with GraphPad Prism using ordinary one-way, two-way ANOVA multiple comparisons tests, or Mann-Whitney t-test. Compiled data is shown as mean±SEM. *p<0.0332, **p<0.0021, ***p<0.0002, ****p<0.0001.

