## Supplemental figures for "Murine cytomegalovirus downregulates ERAAP & induces an unconventional T cell response to self"

**Supplemental Figure 1: ERAAP downregulation in MCMV infected RAW macrophages**

**Supplemental Figure 2: ERAAP downregulation in MCMV infected L cells**

**Supplemental Figure 3: Qa-1b levels remain unchanged in MCMV infected L cells**

Supplemental Figure 1

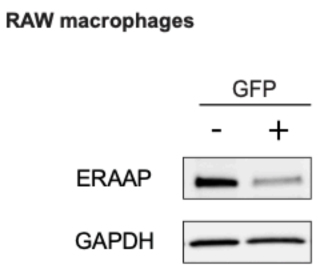


Supplemental Figure 2

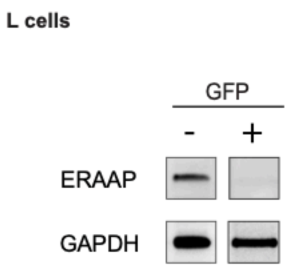


Supplemental Figure 3

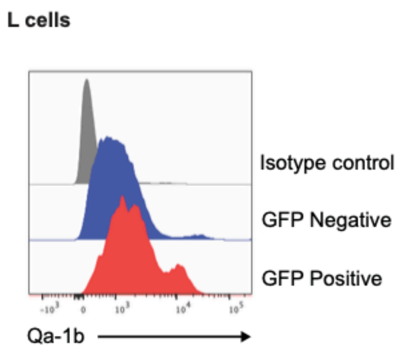


**Supplemental Figure 4: Model for ERAAP downregulation and QFL activation during MCMV infection**

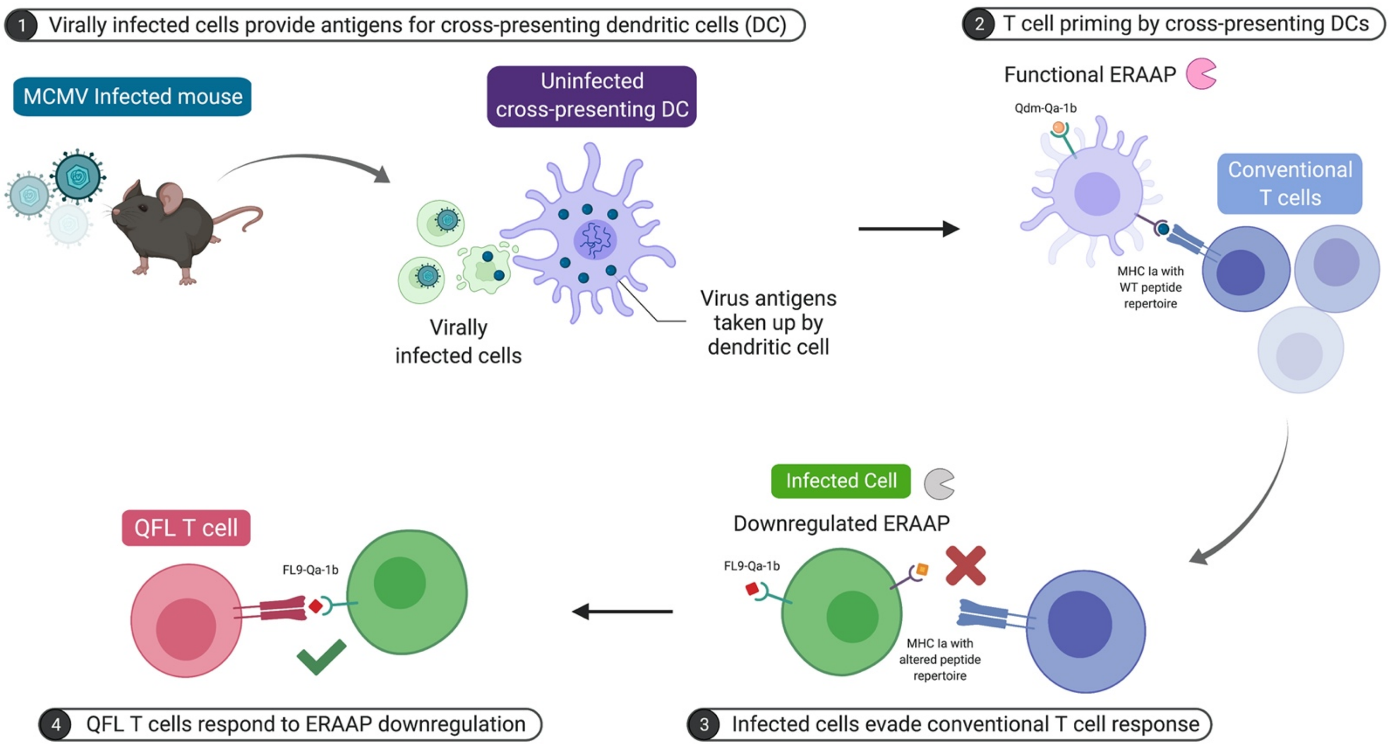
